# Accumulation of ammonium owing to the metabolic imbalance of carbon and nitrogen might inhibit the central metabolism in *Methylomonas* sp. ZR1

**DOI:** 10.1101/2020.01.22.916338

**Authors:** Wei Guo, Ronglin He, Yujie Zhao, Demao Li

## Abstract

The metabolic intermediates of nitrogen source have been proved to have multiple functions on the metabolism of mehthanotrophs. In this study, accumulation and assimilation mechanism of the nitrate metabolic intermediate ammonium in the fast growing *Methylomonas* sp. ZR1 was analyzed. Although, nitrate salt was the best nitrogen source supporting the growth of ZR1, its metabolic intermediate ammonium would accumulate and inhibit ZR1. Kinetic studies indicated that accumulation of NH4^+^ was deduced from the imbalance of nitrogen and carbon metabolism. Compensation of carbon skeleton α-keto-glutaramate could effectively relieve the inhibition of NH4^+^ to ZR1, which further approved the assumption. qPCR analysis indicated a third ammonium assimilation pathway Glycine synthesis system may function in ZR1 under high ammonium tension. In the presence of ammonium, ZR1 might employ two strategies to relieve the ammonium stress, one was assimilating the excess ammonium, and another one was cutting off the nitrogen reduction reactions. Investigation of the nitrogen metabolism and its influence to the carbon metabolism is meaningful to systematically understand and control the C1 feedstock bioconversion process in methanotrophs.

**Importance:** The nitrogen metabolism in methanotrophs has long been concerned. However, there are lots of research problems yet to be solved. In this study, the accumulation and assimilation mechanism of the nitrogen metabolic intermediate ammonium in the fast growing *Methylomonas* sp. ZR1 was analyzed. Owing to the imbalance metabolism of carbon and nitrogen source, ammonium would accumulate to high concentrations to inhibit cell growth. Compensation of carbon skeleton was an effective strategy to relieve the inhibition of NH_4_^+^. A third ammonium assimilation pathway related genes were proved actively expressing in ZR1 when it confronted with high ammonium tension. When confronted with ammonium tension, ZR1 might employ different strategies to relieve the ammonium stress according to the edible carbon source. Revealing the endogenous ammonium accumulation mechanism and its metabolic adjustment effect on the central metabolism of methanotrophs, was meaningful to reveal the complex coordination metabolic mechanism of nitrogen and carbon in methanotrophs.

## Introduction

Methanotrophs are excellent C1 compounds assimilating microorganisms, and its industrial application has a long history since 1960s. At the beginning methanotrophs were used to produce single cell protein from methane or methanol. Nowadays, with the development of biotechnology, methanotrophs were genetic modified to produce lactic acid (Henard et al., 2016; Garg et al., 2018), astaxanthin (Ye and Kelly, 2012) and α-hemelune (Sonntag et al., 2015) from methane. However, the slow growth nature of methanotrophs has constrained its application in industrial biotechnology.

Efforts to optimize fermentation technologies to improve the growth rates of methanotrophs on methane have been reported over the past years (Yu et al., 2003; Park et al., 1991b; Shah et al., 1996; Xing et al., 2006; Han et al., 2009) (Myung et al., 2016). Other factors such as the addition of citrate acid (Xing et al., 2006) and adjusting the ion concentration (Leak and Dalton, 1986) were also reported to have effect in improving the growth rate of methanotrophs from methane. These studies indicated that the growth rate of methanotrophs might be result from multi-factor combined effects. Many studies have also proved that nitrogen sources have strong effect on the growth of methanotrophs. Park et al. studied several factors including the nitrogen source affected the growth of OB3b, and found that nitrate depletion was responsible for the diauxic growth pattern in the batch cultivation of OB3b in the bioreactor. However, its growth declined much with 40 mM nitrate (Park et al., 1991). Hoefman et al. found niche partitioning among methanotrophic species, with methane oxidation activity responses to changes in nitrogen content being dependent on the in situ methanotrophic community structure (Hoefman et al., 2014). Strains have developed a complex mechanism to balance the carbon and nitrogen metabolism (Commichau et al., 2006). Recent studies have shown that many rapidly proliferating cells are dependent on the nitrogen metabolic intermediate such as serine and glutamine (Yang and Vousden, 2016). These studies indicated that nitrogen and its metabolic intermediate might have multiple functions on the central metabolism of methanotrophs, and subsequently affected its growth.

In methanotrophs, the nitrogen metabolism has long been concerned. 1983, Dalton et al. studied ammonia assimilation in the type X methanotroph *Methylococcus capsulatus* Bath, type I methanotroph, *Methylomonas methanica* S1, and the type II methanotroph, *Methylosinus trichosporium* OB3b, and found that, Bath and S1 possess both the glutamine synthase/ glutamine 2-oxo-glutarate amino transferase (GS-GAGOT) and alanine dehydrogenase (ALAD) pathways for the assimilation of ammonia, but operated according to the nitrogen source (Murrellt and Dalton, 1983). Loginova NV et al. (Loginova et al., 1982) studied enzymes involved in ammonium assimilation by 15 bacterial strains of different taxonomy, and found that bacteria were found to differ in the enzymes for ammonium assimilation according to the pathways of primary C1-metabolism. Nyerges (Nyerges, 2008) assessed the differerences in ammonia co-metabolism among four methanotrophs isolates, found the investigated strains exhibited different levels of ammonia and hydroxylamine oxidation, and inhibition of methane-oxidizing activity by ammonia and nitrite. Dam et al. found that ammonium induces differential expression pattern of methane and nitrogen metabolism-related gene in *Methylocystis* sp. strain SC2 (Dam et al., 2014). Kits et al. found that in *Methylomonas denitrifican*, sp. nov. type strain FJG1, methane oxidation could couple to nitrate reduction under hypoxia (Kits et al., 2015). These results revealed that nitrogen metabolism might play an important role in the global metabolism of methanotrophs, and nitrogen metabolism mechanism might be distinct among methanotrophs species.

In this study, the nitrogen metabolism during the cell growth of *Methylomonas* sp. ZR1 were studied at kinetic and gene expression level. It was found that, ZR1 might employ different nitrogen metabolic pattern according to the ediable carbon source, the nitrogen metabolic intermediate ammonium which was also identified as a growth inhibiter was found to accumulate when nitrogen sources were relative surplus in comparison with carbon sources. Carbon skeleton supplementation was found to be an efficient strategy to relieve the inhibition effect of ammonium on the growth of ZR1. qPCR analysis of the carbon and nitrogen metabolic key gene indicated the gene expression diversity when ZR1 was under the ammonium tension condition, and a third ammonia assimilation pathway were found highly expressed with methane as carbon source. All these results further indicated that ammonium might have multidimensional effect on the central metabolism of ZR1.

## Material and Methods

### Strains and culture method

*Methylomonas* sp. ZR1 was isolated by our group and deposited in China General Microbiological Culture Collection Center with the accession number CGMCC No. 9873. It can be cultivated using liquid or solid mineral medium with methane or methanol as the growth substrate. The medium used for ZR1 cultivation was nitrate mineral salts (NMS) medium (Whittenbury et al., 1970). For nitrogen source screening, NMS medium without nitrate salt was added with 1 g/l KNO_3_, 1 g/l NaNO_3_, 0.5 g/l NH_4_Cl, 0.5 g/l (NH_4_)SO_4_, 0.5 g/l urea, 1 g/l trypton or 1 g/l yeast extract separately. Samples having no nitrogen substate were performed as control. To test the ammonium inhibition effect, ammonium chloride at different concentration were added into the NMS medium with or without 1 g/l KNO_3_. And for carbon skeleton compensation test, 0.1 g/l ammonium chloride with 0.3 g/l α-ketoglutaramate (α-KG), 0.4 g/l glutamate, 0.3g/l malic acid, or 0.3g/l pyruvate were simultaneously added into the NMS medium. ZR1 was cultured in flask or bubble column reactor according to the method described by Guo et al. (Guo et al., 2017). When using methanol as carbon source, the initial methanol concentration was 6 g/l without specific instruction. When using methane as carbon source, ZR1 was cultured in bubble column reactor for the ammonium accumulation study; and cultured in flask with gas refreshing every 12 hours for the nitrogen source screening, ammonium inhibition test and carbon skeleton replenishing study.

### Total nitrogen and ammonium concentration analysis method

Total nitrogen concentration in the fermentation broth was analyzed using the TOC/TN analyzer Multi N/C 2100s (Analytik Jena AG, German). Fermentation broth was first centrifuged, and the supernatant were diluted using ddH_2_O into suitable concentration for the analysis.

Ammonium was analyzed using the indophenol blue reaction according to the method described by Xie et al. (Xie et al., 2005). The fermentation broth was first centrifuged, and 0.1 ml of the supernatant were added with 0.5ml reaction solution 1 (3.5 g phenol and 0.04 g sodium nitroprusside in 100 ml ddH_2_O), and 0.5 ml reaction solution 2 (1.8 g sodium sodium hydroxide and 4.0 mmol sodium hypochlorite in 100 ml ddH_2_O). The mixture was maintained at 37°C for 1 hour, and the absorbance of the solutions was read on a spectrophotometer at 625 nm. The concentration of the ammonium in the samples was calculated according to the calibration curve established using ammonium chloride.

### Methanol concentration

Methanol concentration in the fermentation broth were measured by a GC (GC9790, Fuli Instrument, China) equipped with a flame ionization detector (FID) and a capillary column (0.25μm, 60m×0.25mm, 7KG-G013-11 Zebron^TM^, Phenomenex).

### Kinetic analysis of the carbon and nitrogen metabolism to cell growth

The growth, carbon and nitrogen assimilation curve were first fitted using the logistic model in Origin 9.0. Then the simulated curve were took derivative with respect to OD_600_ of ZR1 using the mathematic module of origin 9.0.

### qPCR analysis of expression level of the genes concerns carbon and ammonium metabolism

Strain ZR1 was initially grown up to log phase (OD_600_ achieved 0.8) in 50 ml of NMS medium in a 250 ml bottle fitted with butyl rubber septum and with 20% methane (v/v) in the headspace. For methanol as carbon sources, strains were cultured with 6 g/l methanol in 50 ml NMS medium in 250 ml flask. For preparing ammonium inhibition samples, 0.1 g/l of NH_4_Cl were then added into the medium. For carbon compensation samples, 0.1 g/l of NH_4_Cl and 0.3 g/l of α-keto-glutaramate (α-KG) were added, and samples without any operation were performed as control. All samples were incubated for 2 hours, then, cells were collected by centrifugation and washed with TE buffer. RNA was extracted immediately using the RNAprep Pure Cell/Bacteria Kit (Tiangen Biotech (Beijing) CO., Ltd), according to the product instructions. The extracted RNA was used as template to construct cDNA using the Takara Prime Script RTreagent Kit with gDNA Eraser (Perfect Real Time). Then genes were amplified using the SYBR Premix EX Taq kit, using Applied Biosystems 7500 fast Real-Time PCR system. Standard housekeeping gene 16S rDNA was selected as internal control gene. qPCR data obtained were analyzed using the method described by Pfaffl (Pfaffl, 2001).

Primers used are listed in table 1.

**Table 1.**
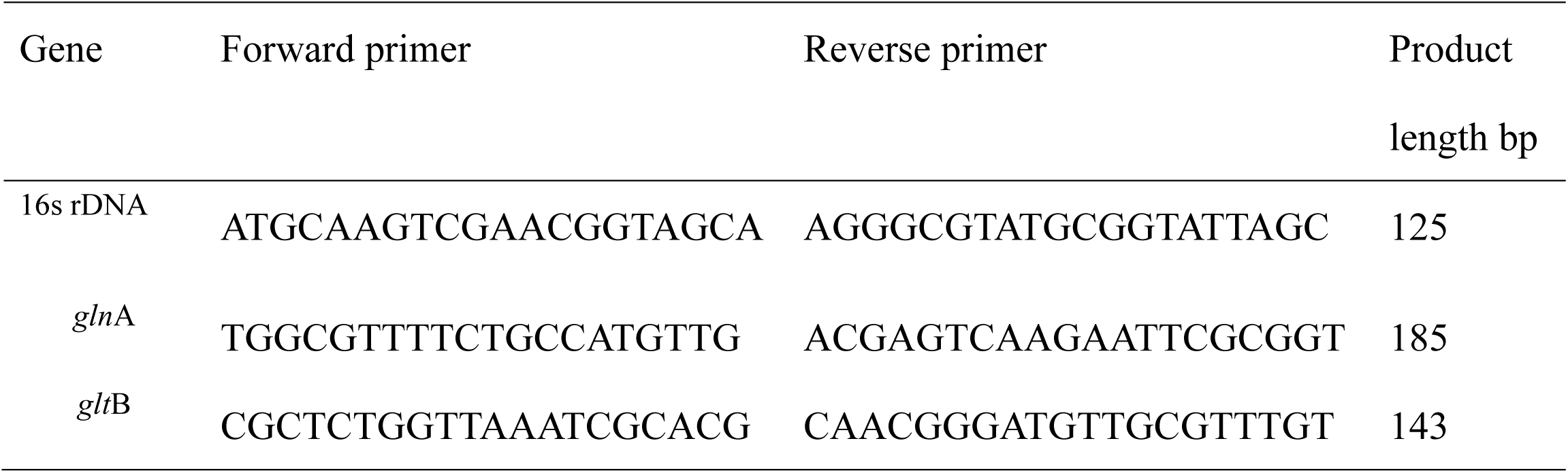
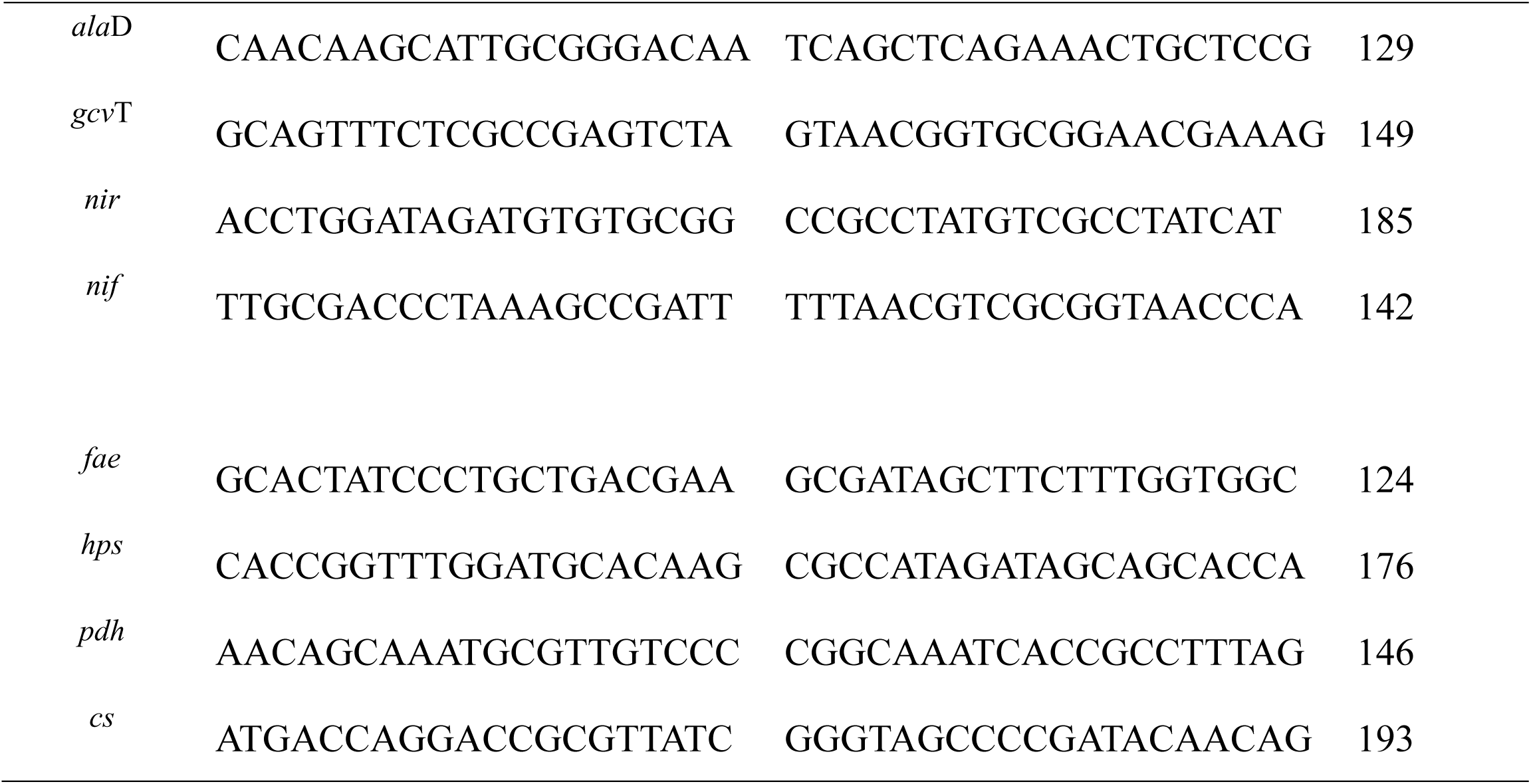
Primers designed for qPCR

All culture conditions were performed in triplicate biological replicates, and for qPCR analysis, each biological sample was carried out in triplicate. Raw data of the qPCR result were shown in supplementary Table S1.

## Result and discussion

### 1. Nitrogen source effect for the growth of ZR1 from methane and methanol

Seven categories of nitrogen substrate were tested for the growth of ZR1 from both methane and methanol (Figure 1). According to Figure 1, nitrate salts were the best nitrogen sources supporting the growth of ZR1 from both methanol and methane among the tested compounds. And the growth of ZR1 from both methanol and methane with ammonium salts was much weaker than from nitrate salts. Meanwhile, the effects of the nitrogen sources on the growth of ZR1 from methane and methanol were somewhat different. With methane as carbon sources the cell density (OD_600_) of ZR1 achieved 2.5 with potassium nitrate, 1.8 with sodium nitrate, 1.8 with urea, 1.5 with yeast extract, 0.5 with tryptone, and 0.34 with ammonia chloride, 0.35 from ammonia sulfate. With methanol as carbon sources the cell density of ZR1 achieved 8.2 from sodium nitrate, 7.9 from potassium nitrate, 3.9 from yeast extract, 1.8 from urea, 0.4 from tryptone, 0.35 from ammonia chloride, and 0.33 from ammonia sulfate. It can be seen that nitrate salt was the most suitable nitrogen source and ammonia salt could not effectively support the growth of ZR1 from methane and methanol.

It was generally regarded that ammonium radicals were the competitive inhibitor of particular methane monooxygenase (pMMO) (He et al., 2017; Hu and Lu, 2015; Dam et al., 2014; Nyerges et al., 2010; Nyerges and Stein, 2009; Nyerges, 2008; Dunfield and Knowles, 1995; Schnell and King, 1994; Carlsen et al., 1991; Murrellt and Dalton, 1983), which will resulted in the lower growth of strains from methane. However, this study also proved that ammonium would hinder the growth of ZR1 from methanol, although its supposed competent object pMMO is not the key enzyme for the metabolism of methanol. It means that ammonium might have other inhibition effects for the growth of ZR1, besides its competitive inhibition effect to pMMO.

**Figure. 1.**
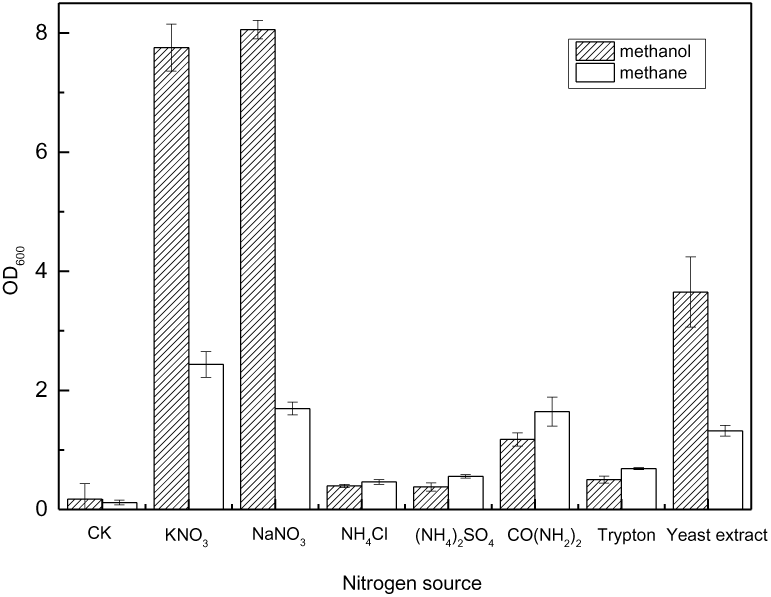
Growth of ZR1 in the presence of different nitrogen substrate from methane and methanol

### 2. Effect of nitrate concentration on the growth of ZR1 from methane and methanol

According to Figure 1, nitrate salts were supposed to be the most suitable nitrate source that supporting the growth of ZR1 from both methane and methanol, and effects of potassium nitrate concentration on the growth of ZR1 were investigated subsequently. According to Fig. 2, the obvious substrate inhibition effects of nitrate salts on the growth of ZR1 were investigated. The highest cell density of ZR1 achieved 7.8 with 1.5 g/l potassium nitrate (Figure 2a). Meanwhile, Figure 2b indicated that the growth rate of ZR1 is first ascend with the increase in the substrate level and approaches a maximum value at 1.5 g/l of potassium nitrate. Then, a subsequent increase in the nitrogen concentration leaded to a decrease in the specific growth rate of ZR1. The situation with methane as carbon sources is similar to that of methanol. The highest cell density of ZR1 achieved 4.8 with 1.5 g/l potassium nitrate (Figure 2c), however the specific growth rate of ZR1 achieved 0.23 with 1 g/l of potassium nitrate (Figure 2d). It has been reported that growth of type Ⅱ methanotrophs OB3b were also inhibited by 40 mM nitrate (4 g/l)(Park et al., 1991a). These phenomenon indicated that nitrate as nitrogen source when its concentration achieved a certain value might inhibit the growth of methanotrophs. According to the substrate inhibition theory (Muchandani and Luong, 1989), an increase in the substrate concentration could cause an alteration in the cell metabolism such as an overproduction of a molecule by one pathway which results in the feedback inhibition of a second related pathway. Thus the nitrate metabolism pathway of ZR1 was further explored.

**Figure 2.**
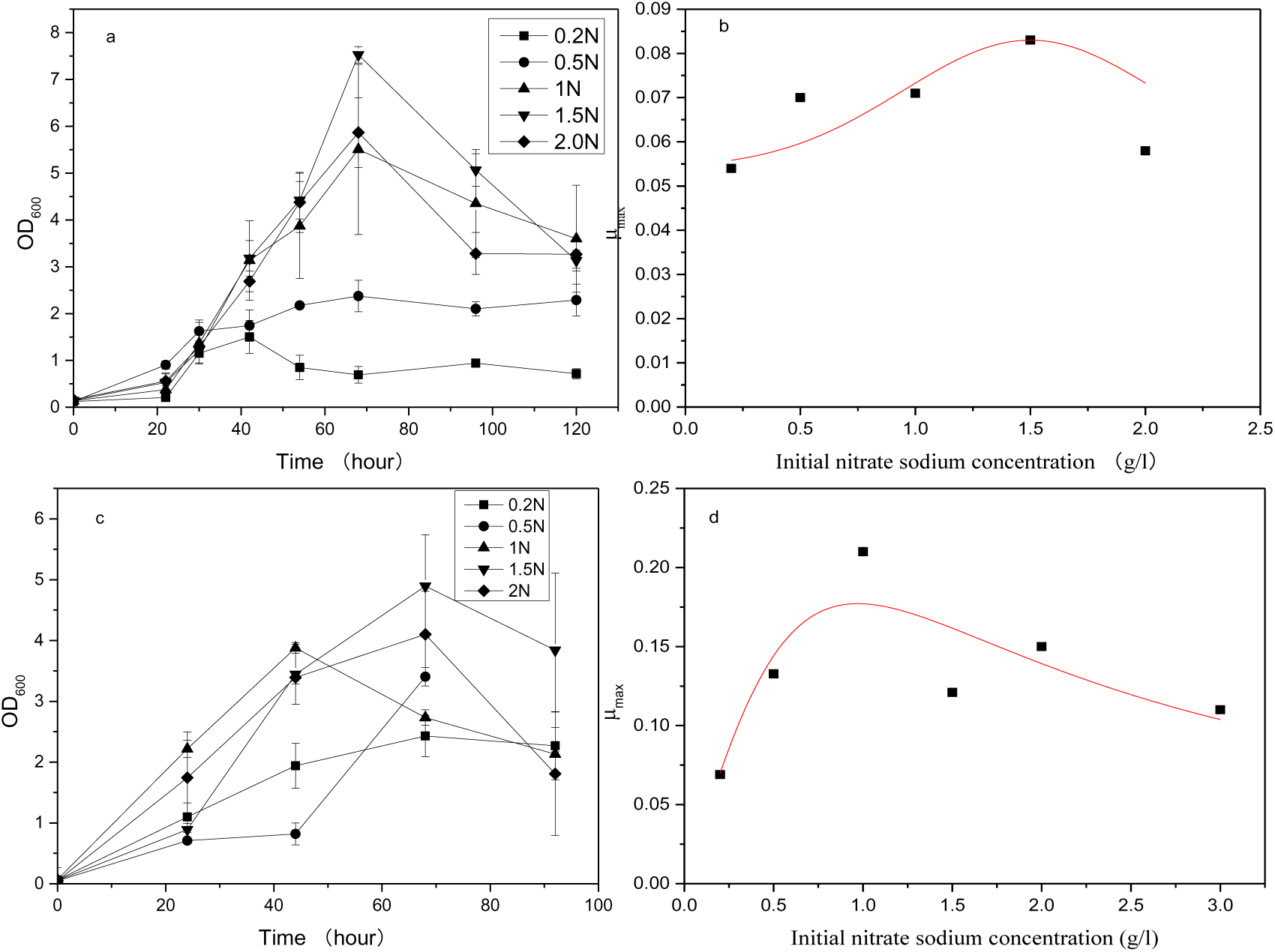
Effect of nitrate concentration on the growth of ZR1 from methane and methanol a, growth of ZR1 with different initial nitrate sodium concentration from methanol; b, specific growth rate of ZR1 with different initial nitrate concentration from methanol; c, growth of ZR1 with different initial nitrate sodium concentration from methane; d, specific growth rate of ZR1 with different initial nitrate concentration from methanol;

### 3. The formulation of ammonium, and its inhibition effect to ZR1

Considering the nitrate metabolism pathway of methanotrophs, nitrogen metabolism intermediate NH_4_^+^ was supposed to be accumulated during the process. Thus ammonium concentration in the fermentation broth was analyzed during the fermentation process of ZR1 from methane and methanol. According to Figure 3, the concentration of NH_4_^+^ in the fermentation broth grew higher and higher during the fermentation process, which indicated that NH_4_^+^ might accumulated accompanied with the growth of ZR1 when excess nitrate were supplied. With the increasing concentration of nitrogen source, the starting time of NH_4_^+^ accumulation has been moved up. Meanwhile the final concentration of NH_4_^+^ increased with the increase of the initial concentration of KNO_3_. With 3 g/l of KNO_3_, the final accumulated NH_4_^+^ achieved 25 mg/l with methanol as carbon sources and 16 mg/l with methane as carbon source. According to Fig. 1, NH_4_^+^ was supposed to be an inhibitor of the growth of ZR1. The accumulated NH_4_^+^ during the growth process of ZR1 might inhibit the growth of ZR1. Thus the effect of NH_4_^+^ on the growth of ZR1 was further analyzed.

**Figure. 3.**
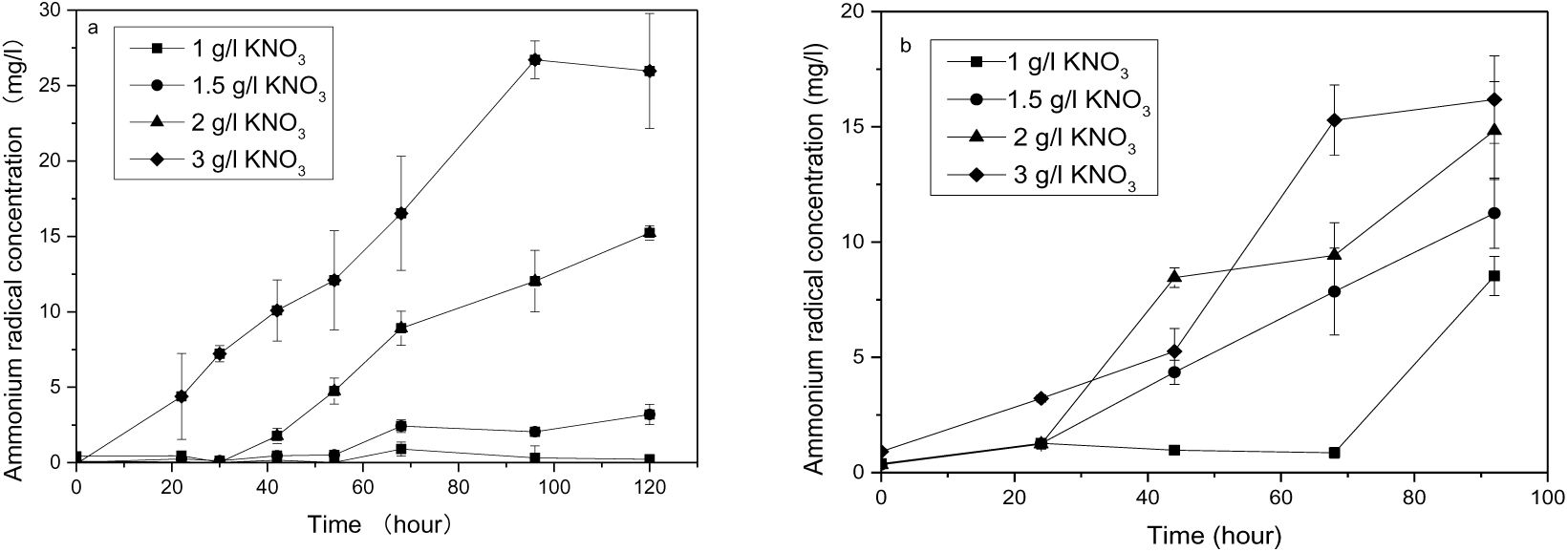
Ammonium accumulation of ZR1 in the fermentation broth with different initiate nitrate concentration from methanol and methane

Growths of ZR1 with NH_4_Cl as sole nitrogen source or with both NH_4_Cl and KNO_3_ at different concentrations were investigated. It was found that, when NH_4_Cl were used as the mono-nitrogen sources, cell growth of ZR1 first rise up with the going up of the NH_4_Cl concentration, the highest cell growth achieved 1.5 when NH_4_Cl achieved 0.05, and then growth of ZR1 from methane decreased when the initial NH_4_Cl concentration growing higher, and OD_600_ of ZR1 with NH_4_Cl at 0.1-0.5 g/l only achieved 0.42, much lower than the control (with 1 g/l KNO_3_ as nitrogen source). When 1 g/l of KNO_3_ with addition of 0.05 g/l of NH_4_Cl was used as nitrogen sources, the highest cell density of ZR1 from methane achieved 2.5, much higher than control. While the highest cell density of ZR1 from methanol only achieved 0.42 under the same condition. When 1 g/l of KNO_3_ with addition of 0.1 g/l of NH_4_Cl (NH_4_^+^ 34 mg/l, 1.86 mmol/l) was used as nitrogen sources, the highest cell density of ZR1 only achieved 0.7 with methane and 0.8 with methanol. These results showed that, with the existence of KNO_3_, growth of ZR1 from methane and methanol was inhibited by NH_4_Cl at concentration higher than 0.1 g/l. Growth of ZR1 from methane could resist with 0.05 g/l of NH_4_Cl (0. 93 mmol/l), and were strictly inhibited with 0.1 g/l of NH_4_Cl (1.86 mmol/l), while growth of ZR1 from methanol were totally inhibited by 0.05 g/l of NH_4_Cl, which indicated that growth of ZR1 from methanol is more sensitive to NH_4_^+^ than that from methane. And according to Figure 3 the final accumulated NH_4_^+^ with initial higher concentration of nitrate salts could achieve 25 mg/l (0.89 mmol/l), which was high enough to inhibit the growth of ZR1 from methanol.

It was generally regarded that ammonium radicals were the competitive inhibitor of pMMO (He et al., 2017). In this study, ammonium was also identified to inhibit the growth of ZR1 from methanol and accumulate during the growing process. Recently, ammonium was reported to have effects on global gene expression of methane and nitrogen metabolism-related gene in methanotrophs (Dam et al., 2014). Being the nitrogen metabolic intermediate, accumulation of ammonium may have multiple effects on the growth of methanotrophs. Revealing its accumulation and assimilation mechanism is meaningful to understand the metabolism mechanism of methanotrophs.

### 4. Carbon and nitrogen metabolic kinetic analysis of ZR1 from methanol

According to Figure 4, concentration of the accumulated NH_4_^+^ was proportional to the initial nitrogen concentration in the fermentation broth. And the accumulation of NH_4_^+^ might be result from the high initial nitrogen source concentration (in another words the lacking of the carbon skeleton). Thus, co-metabolism of the carbon and nitrogen and accumulation of ammonium in ZR1 with methanol as carbon source was further analyzed.

**Figure. 4.**
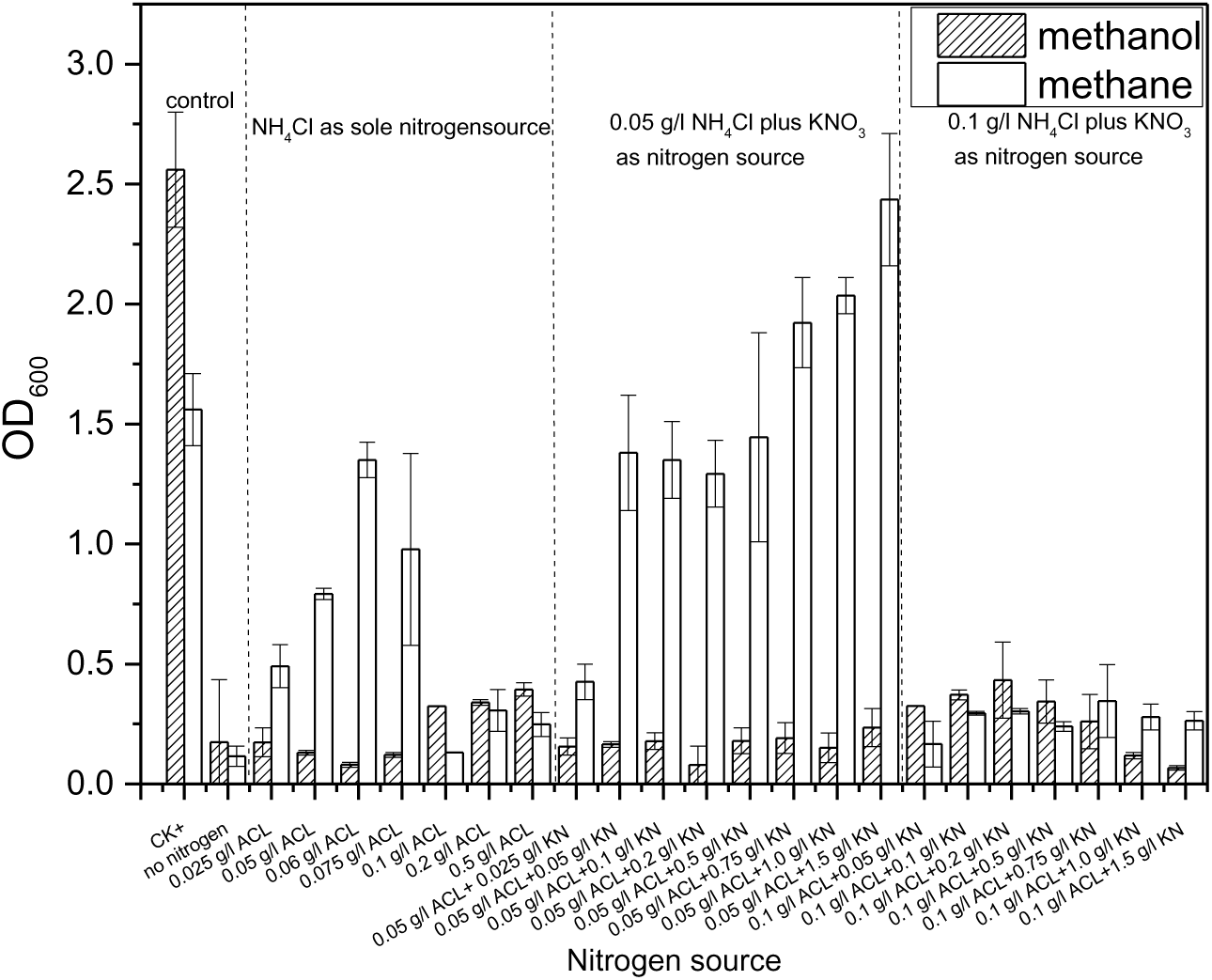
Effect of Ammonium on the growth of ZR1 from methane and methanol. ACL, NH_4_Cl; KN, KNO_3_;

According to Figure 5, it can be found that with methanol under the same concentration of 6 g/l, ammonium accumulation increased with the increscent of nitrogen concentration. Nonetheless, increasing the carbon sources to 12 g/l (Figure 5e), the accumulated ammonium will subsequently decrease. These results indicated that ammonium accumulation was deduced from the imbalance of the carbon and nitrogen metabolism. And according to the nitrogen test result (Figure 1, Figure 3), the accumulated ammonium could achieve 25-30 g/l which was high enough to inhibit the growth of ZR1 from methane and methanol as carbon sources.

**Figure 5.**
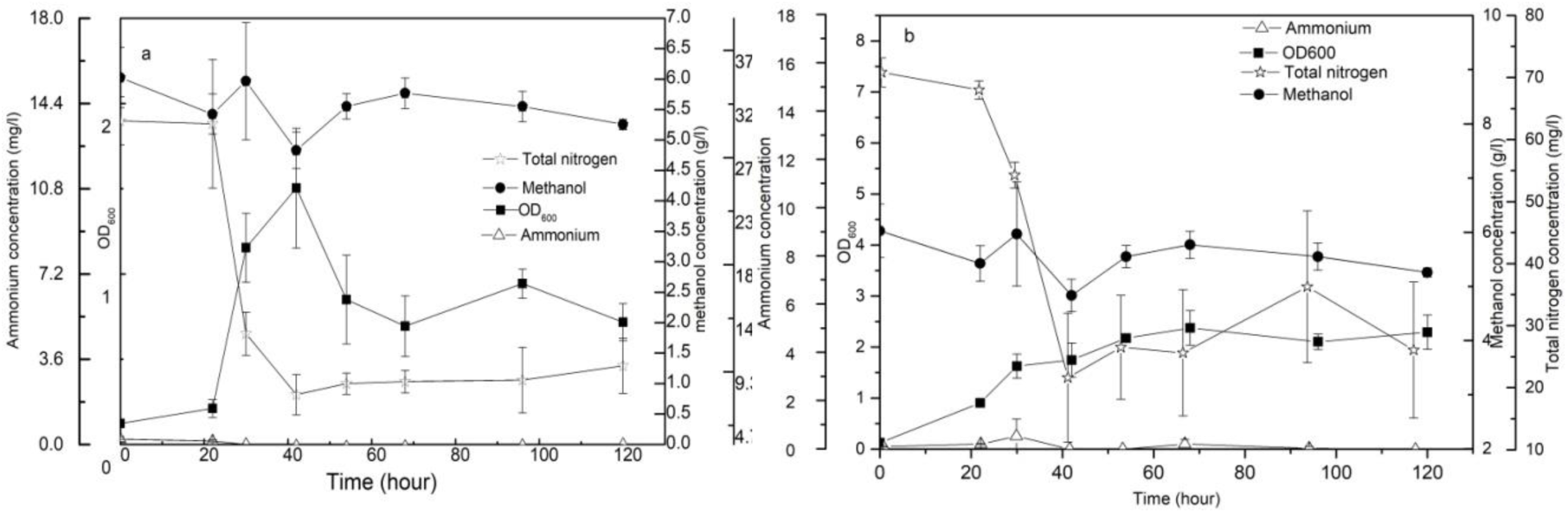
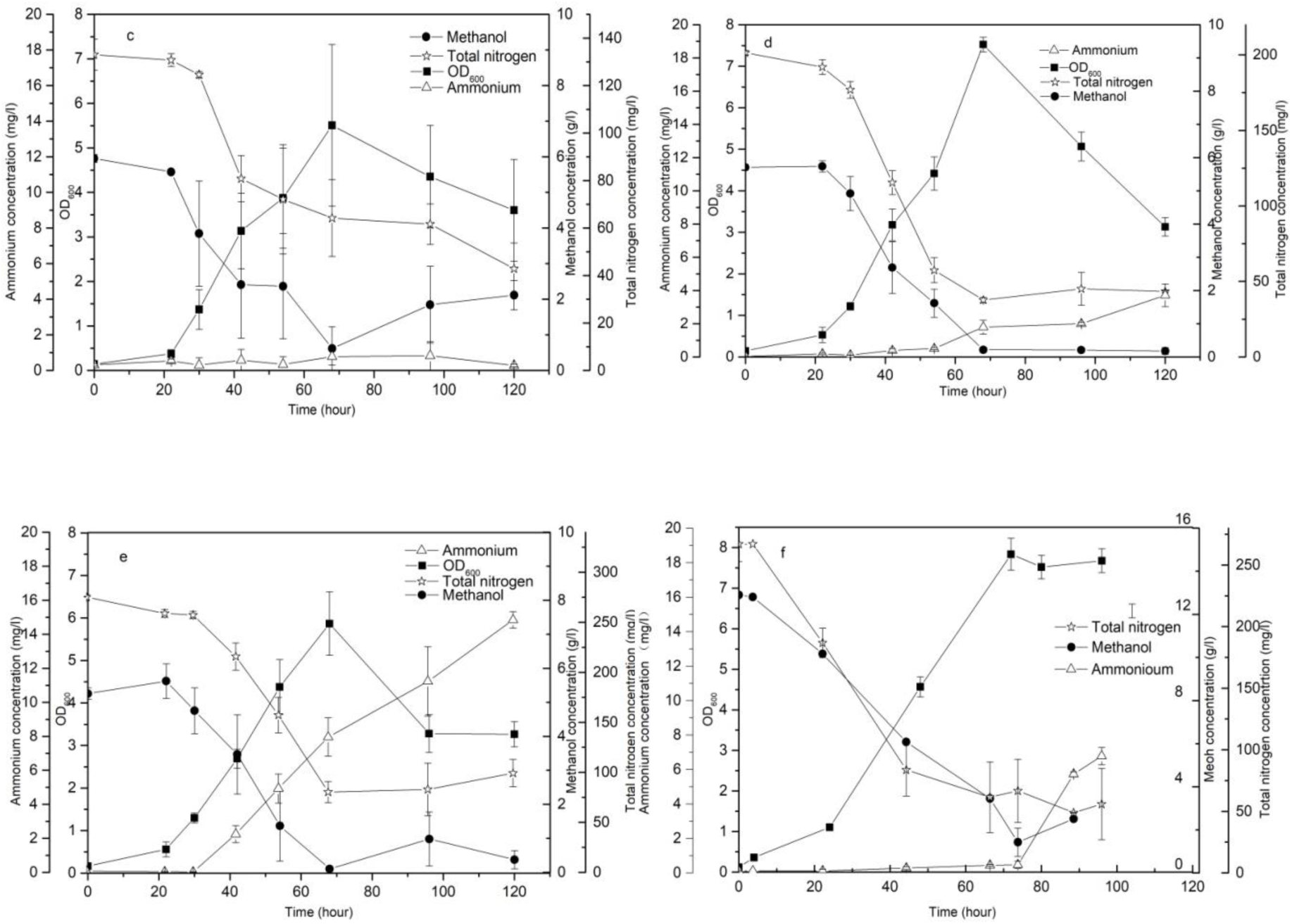
Kinetic analysis of the carbon and nitrogen metabolism of ZR1 from methanol. a, 6 g/l methanol and 0.2 g/l nitrate sodium; b.6g/l methanol and 0.5g/l nitrate sodium; c. 6 g/l methanol and 1 g/l nitrate sodium; d. 6 g/l methanol and 1.5 g/l nitrate sodium; e. 6 g/l methanol and 2 g/l nitrate sodium; f. 12g/l methanol and 2g/l nitrate sodium

To further compare the carbon and nitrogen metabolism pattern, specific consuming rate of the carbon and nitrogen during the growth process of ZR1 were further analyzed. According to Figure 6, it can be seen that the specific uptake rates of nitrogen and carbon dynamically changed with the nitrogen-carbon ratio (NCR). High NCR will result in delayed carbon consuming, and the carbon and nitrogen uptake will keep in step when NCR fall into a suitable value.

**Figure 6.**
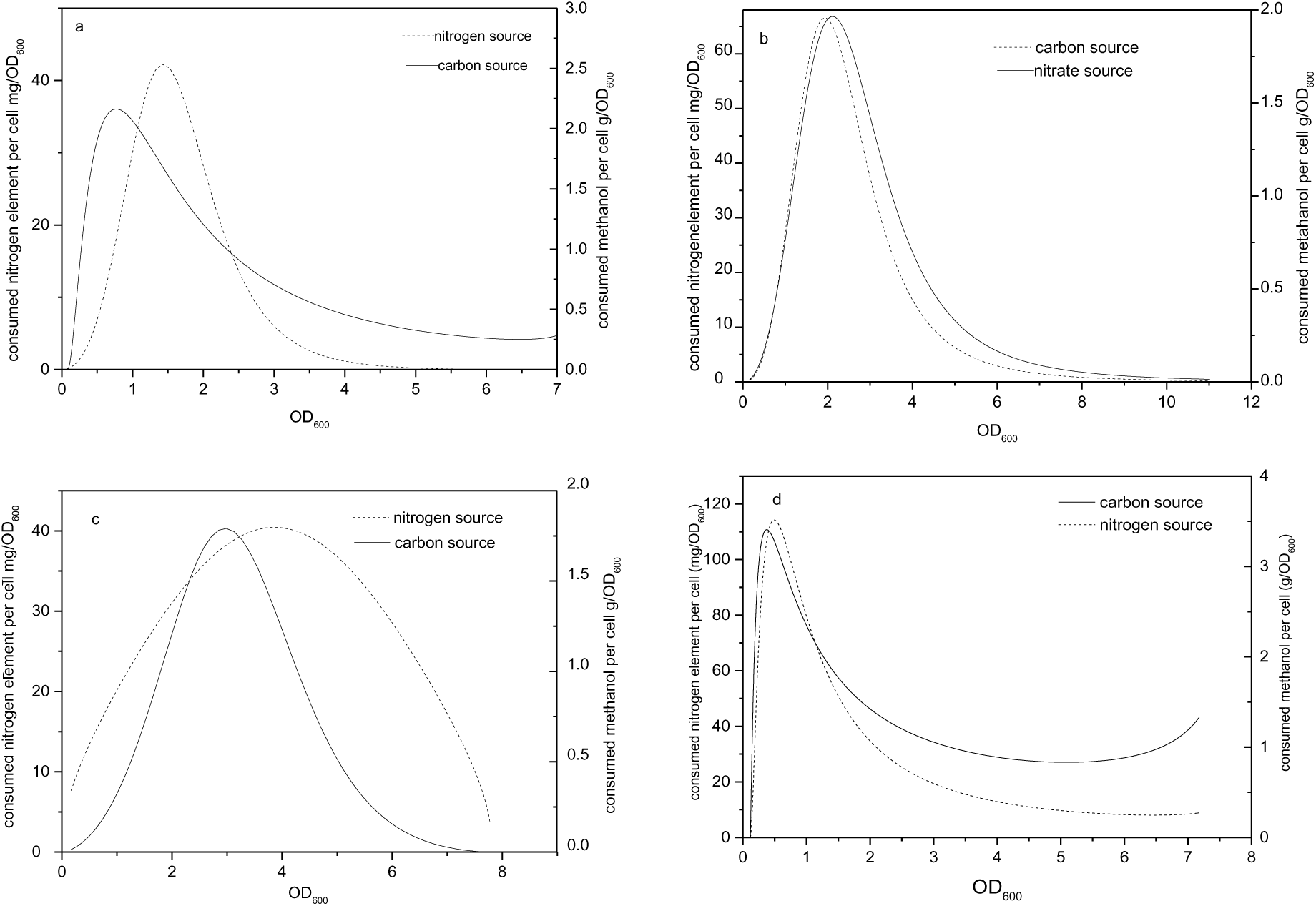
Kinetic analysis of the carbon and nitrogen metabolism to cell growth a. 6 g/l methanol and 1g/l nitrate sodium; b.6g/l methanol and 1.5g/l nitrate sodium; c. 6 g/l methanol and 2 g/l nitrate sodium; d. 12g/l methanol and 2g/l nitrate sodium

### 5. Relieving of ammonium inhibition effect by the carbon metabolites

According to the nitrogen sources metabolic pathway (Murrellt and Dalton, 1983; He et al., 2017; Nyerges, 2008a), methanotrophs mainly assimilate ammonium through the glutamine synthase/ glutamine 2-oxo-glutarate amino transferase (GS-GAGOT) and alanine dehydrogenase (ALAD) pathways. To further confirm that accumulation of NH_4_^+^ was derived from the shortness of the carbon skeleton, several carbon metabolic mediates pyruvate, malic acid, citrate acid, α-KG which are related with the nitrogen metabolism were added to the ammonium accumulation samples (Figure. 7). According to Figure 7, α -KG and glutamate were the most effective carbon metabolic mediates in relieving of the NH_4_^+^ inhibition effect with methane as carbon sources. However, with methanol as carbon source only α-KG could effectively relieve the inhibition effect of NH_4_^+^ on the growth of ZR1. Carbon metabolic mediates upstream or downstream of α -KG, has small relieving effect on NH_4_^+^ inhibition. It was also found that, direct supply of carbon skeleton such as pyruvate, citric acid, α-KG, glutamate or malic acid have somewhat inhibit effect on the growth of ZR1 from methanol. Meanwhile,α-KG and glutamate were found to have stimulate effect on the growth of ZR1 from methane.

**Figure 7.**
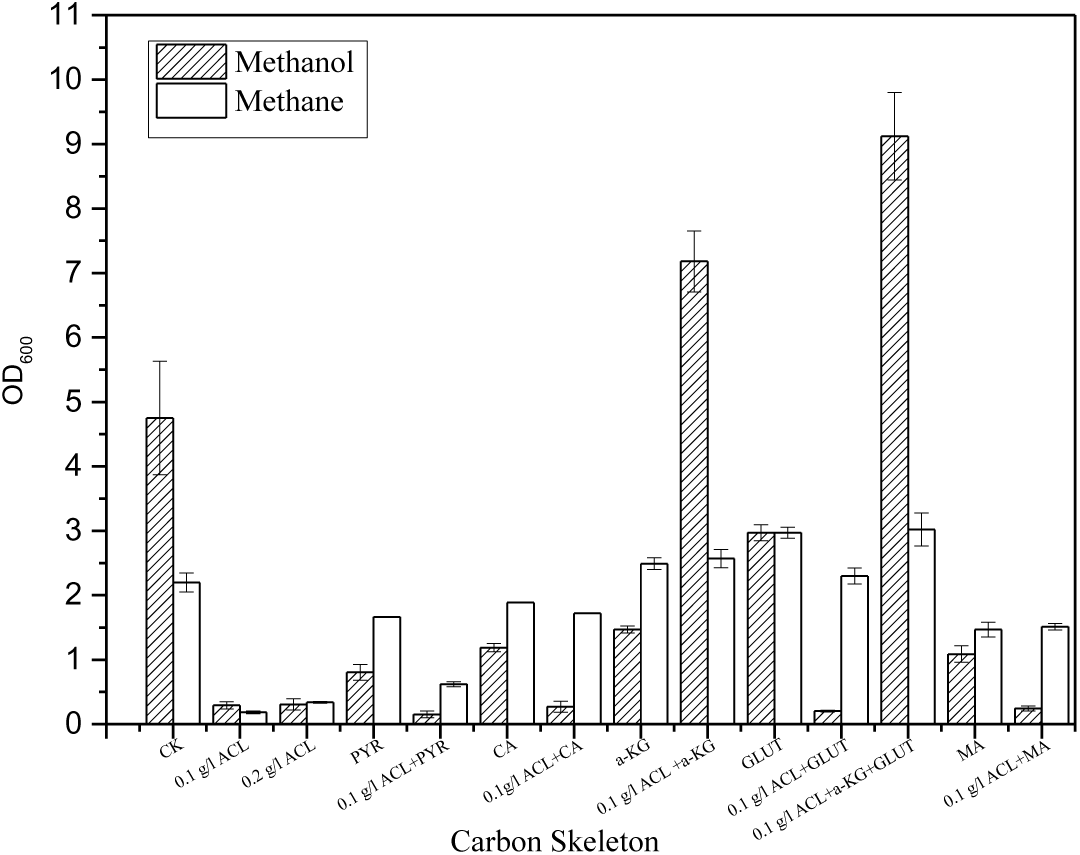
Relieve Effect of the carbon skeleton to ammonium inhibition CK: control; ACL, NH_4_Cl; PYR, pyruvate; CA, citrate acid; α-KG, α -ketoglutaramate; GLUT, glutamate; MA, malic acid;

It means that, ZR1 may utilize different carbon and nitrogen metabolic mechanism to balance the carbon and nitrogen metabolism. Nonetheless, α-KG play more important role in assimilation of the nitrogen metabolic intermediate ammonium.

### 6. Transcript level analysis the carbon and nitrogen metabolism in ZR1 with the accumulation of ammonium

According to the nitrogen metabolism pathway of microorganisms, many studies indicated that, ammonia is a competitive inhibitor of pMMO, which deduced the inhibition effect of methanotrophs from methane. However, this study also identified the inhibition effect of NH_4_^+^ on ZR1 with methanol as carbon source. So, besides the competitive inhibition effect of MMO, NH_4_^+^ as a nitrogen metabolism intermediate may have multiple complex effects on the metabolic pathway of methanotrophs. Thus transcript level analysis of the genes relevant with carbon and nitrogen metabolism in ZR1 was carried out by qPCR.

Ammonium mainly transferred to the C skeletons of pyruvate or intermediates of citric acid cycle (glycogenic) and others to form amino acid. Most amino acid participate in transamination reactions with TCA cycle mediate such as oxaloacetate, or α-ketoglutarate to form aspartate, or glutamate, respectively, and the α-KG corresponding to the original ammonium assimilation. Based on the genomic analysis result of ZR1, besides the ALAD and GS/GOGAT pathway, another special ammonium assimilation mechanism the Glycine synthase system may exist and function in methanotrphs(Figure 8). In this study, the GS-GAGOT, ALAD, and Glycine synthesis system on behalf of three main pathways of the ammonium assimilation were studied by qPCRT, the data quality of the qPCR result was presented in supplementary Table 1. E-supplementary data of this work can be found in online version of the paper. The relative expression fold of the target genes was listed in Table 2.

**Figure 8.**
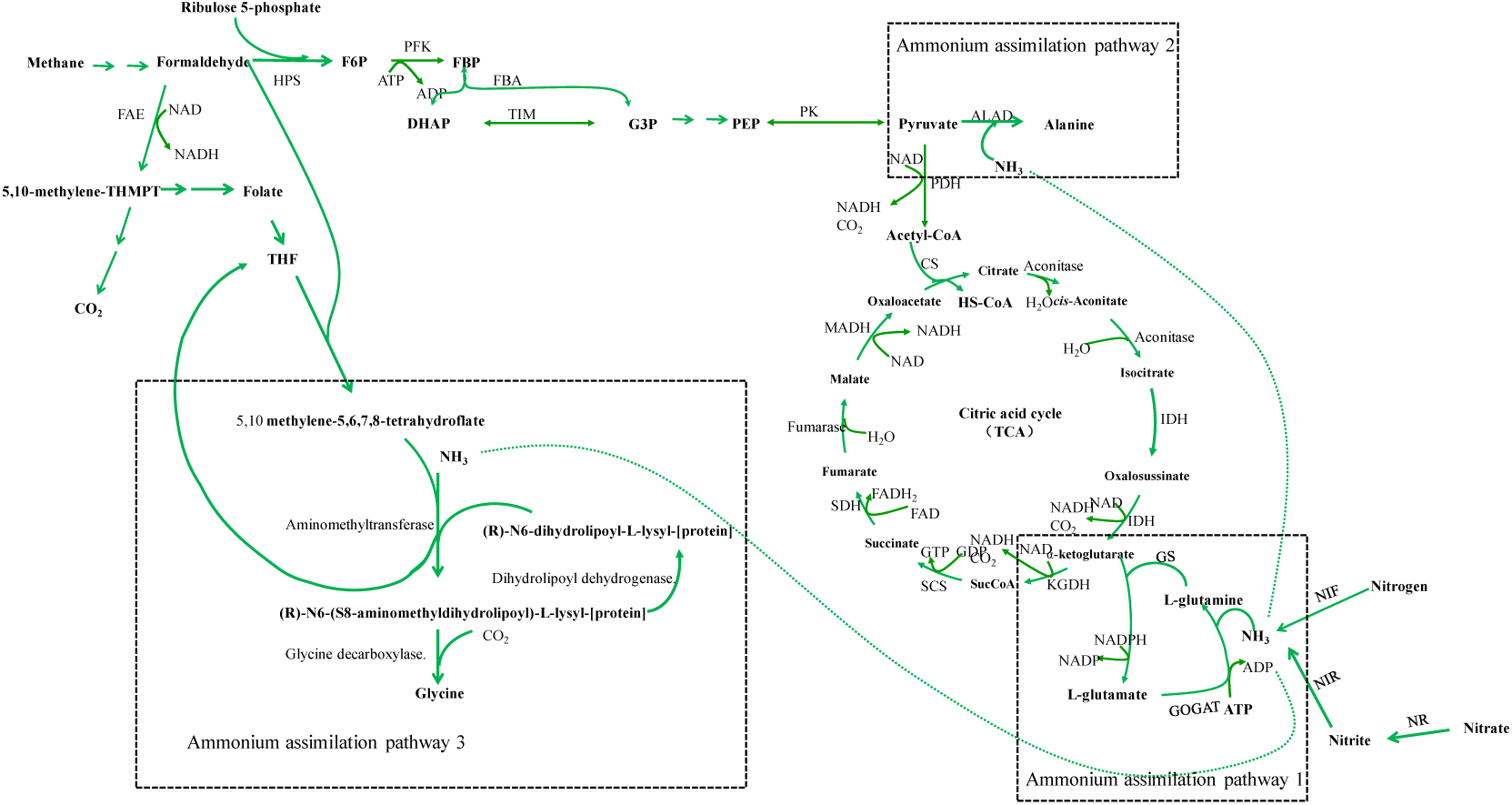
Nitrogen metabolism pathway of ZR1

**Table 2.** Relative expression level of the genes concerning nitrogen and carbon metabolism

| Mean fold change in gene expression of samples |  |  |  |  |  |  |  |  |
| --- | --- | --- | --- | --- | --- | --- | --- | --- |
| genes | protein | N<sub>control</sub> | N<sub>ACL</sub> | N<sub>NCL-KG</sub> | M<sub>control</sub>-N<sub>control</sub> | M<sub>control</sub> | M<sub>ACL</sub> | M<sub>ACL-KG</sub> |
| *glnA* | Glutamine synthase | 1 | 1.92 | 15.73 | 0.69 | 1.00 | 0.89 | 1.34 |
| *gltB* | Glutamate synthase large subunit | 1 | 7.48 | 12.64 | 42.38 | 1.00 | 0.45 | 0.12 |
| *alaD* | Alanine dehydrogenase | 1 | 30.84 | 17.84 | 25.07 | 1.00 | 0.48 | 0.08 |
| *gcvT* | Aminomethyltransferase | 1 | 19.08 | 2.96 | 1.56 | 1.00 | 0.95 | 2.17 |
| *nir* | Nitrite reductase | 1 | 8.74 | 1.93 | 2.01 | 1.00 | 0.48 | 0.92 |
| *nif* | Nitrogenase | 1 | 4.59 | 4.14 | 2.15 | 1.00 | 0.50 | 0.92 |
| *fae* | Formaldehyde activating enzyme | 1 | 4.32 | 2.94 | 2.99 | 1.00 | 1.07 | 1.04 |
| *hps* | 3-hexulose-6-phosphate synthase | 1 | 13.30 | 2.53 | 5.70 | 1.00 | 1.42 | 0.75 |
| *pdh* | Pyruvate Dehydrogenase | 1 | 1.66 | 1.35 | 0.05 | 1.00 | 1.88 | 2.06 |
| *cs* | Citrate synthase | 1 | 1.15 | 1.52 | 8.32 | 1.00 | 1.24 | 1.21 |

HPS:3-hexulose-6-phosphate synthase, DHAP, Dihydroxyacetone phosphate; F6P, Fructose-6-phosphate; FBP, Fructose-1, 6 –bisphosphate; G3P, Glyceraldehyde 3-phosphate; PEP, Phosphoenolopyruvate; SucCoA, Succinyl-CoA; TIM, Triosephosphate isomerase; GAPD, Glyceraldehyde 3-phosphate dehydrogenase; PK, pyruvate kinase; PFK, Phosphofructose kinase; CS, Citrate synthase; IDH, Isocitrate dehydrogenase; KGDH, Ketoglutarate dehydrogenase; SCS, Succinyl-CoA synthetase; SDH, Succinate dehydrogenase; MDH, Malate dehydrogenase; GS, Glutamine synthetase; GOGAT, Glutamate synthase; cd1 NIRs, Cytochrome cd1 nitrite reductase; ALAD, Alanine dehydrogenase; NIF, Nitrogenase; FAE, Formaldehyde activating enzyme;

Nitrogen assimilation pathway 1:

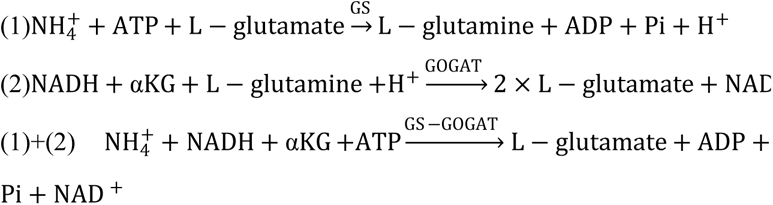

Nitrogen assimilation pathway 2:

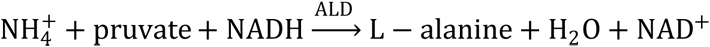

Nitrogen assimilation pathway 3:

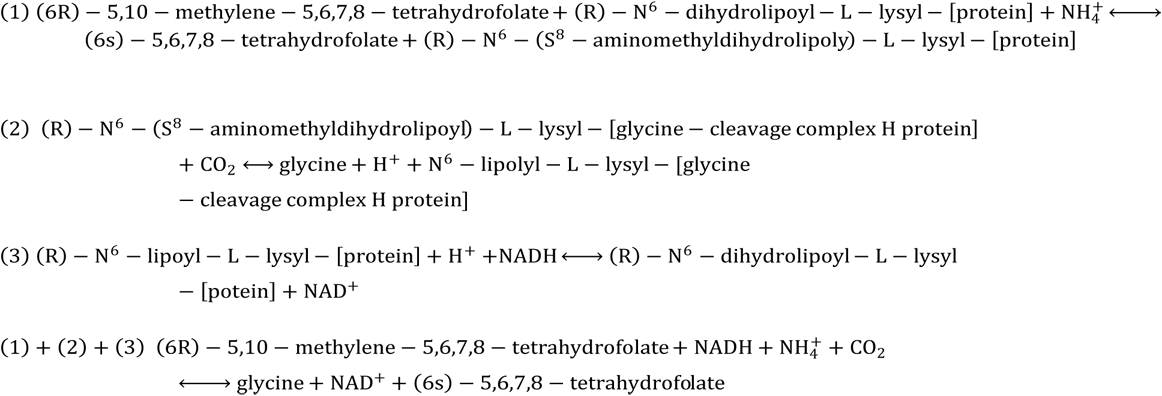

Direction 1 is catalyzed by two enzymes, glutamine synthase and glutamine 2-oxo-glutarate amino transferase. Direction 2 is catalyzed by the alanine dehydrogenase. Direction 3 is catalyzed by the Glycine synthesis system.

N_control_, Samples cultured with methane; N_ACL_, Samples cultured with methane under 0.1 g/l NH_4_Cl condition; N_ACL-KG_, Samples cultured with methane under 0.1 g/l NH_4_Cl and 0.3 g/l α-KG; Mcontrol, Samples cultured with methanol; M_ACL_, Samples cultured with methanol under 0.1 g/l NH_4_Cl condition; M_ACL-KG_, Samples cultured with methanol under 0.1 g/l NH_4_Cl and 0.3 g/l α-KG;

According to Table 2, with methane as carbon source, when 0.1 g/l ammonium accumulated, the *ala*D and *gcv*T gene expression levels were 30.84 and 19.08 times higher than the control, while the genes *gln*A and *glt*B which responsible for the assimilation of NH_4_^+^ with the C5 substrate expressed 1.92 and 7.48 times higher than that of the control. This indicated that ZR1 may rely more on the C3 carbon skeleton to assimilate the extra NH_4_^+^. Another interesting phenomenon is the high expression level of *gcv*T gene (19.08) when ZR1 confronted with 0.1g/l of NH_4_Cl with methane as carbon source and its expression level returned to 2.96 when the C5 carbon skeleton was replenished. According the third direction of nitrogen assimilation, this reaction consumes much 5,10-methylene-5,6,7,8-tetrahydrofolate which is a derivative of C1 substrate. And the exhaustion of the C1 and C3 substrate may reduce the flux distribution of the downstream carbon metabolic pathway and cause the worse growth of ZR1 when high concentration ammonium appeared.

Furthermore, NH_4_^+^ accumulation may also induce the expression of the *hps* gene which is responsible for the assimilation of C1 substrate. Surprisingly, expression level of genes concerns the nitrogen assimilation and formaldehyde oxidization was also up-regulated 4-8 times. On the other hand, with methanol as carbon sources, ZR1 possess a relative high expression background with the *glt*B, *ala*D, *hps* and *cs* genes. And genes changed slightly with NH_4_^+^ accumulation in comparison with control. However, if considering the relative gene expression level of the samples with methanol as carbon sources in comparison with methane, the C5 and C3 carbon skeleton assimilation gene *glt*B and *ala*D of ZR1 still kept 19.07 and 12.03 times higher than the methane control. Thus, when confronted with high ammonium concentration, ZR1 may mainly assimilate NH_4_^+^ through pathway 1 and 2. Meanwhile, decreasing the *nir* and *nif* gene expression level to decrease the formulation of ammonium might be the main strategy employed by ZR1. With the replenishment of the C5 skeleton, the expression level of ammonium assimilation gene *glt*B and *ala*D using C5 and C3 carbon skeleton decreased to 0.12 and 0.08, and the nitrate and nitrogen reduction gene return to normal level. However, the *gcv*T gene in the third ammonium assimilation direction expressed 2 times higher.

Nitrate and its metabolic intermediate were found to have multiple functions for the primary metabolism in biology (Stitt, 1999; Commichau et al., 2006; Cueto et al., 2016). Ammonium was found to be an important signal molecular to affect the methane and nitrogen metabolism in methanotrophs (Dam et al., 2014; Bodelier and Laanbroek, 2004). In this study, it was also found that, owing to the imbalance metabolism of carbon and nitrogen source, ammonium would accumulated to congcentrations high enough to inhibited cell growth. High concentration ammonium will result in high level expression of several genes concerned the carbon and nitrogen metabolism.

## Conclusions

Nitrate and its metabolic intermediate were found to have multiple functions for the primary metabolism in biology (Stitt, 1999; Commichau et al., 2006; Cueto-Rojas et al., 2016). Ammonium was found to be an important signal molecular to affect the methane and nitrogen metabolism in methanotrophs (Dam et al., 2014; Bodelier and Laanbroek, 2004). In this study, it was also found that, owing to the imbalance metabolism of carbon and nitrogen source, ammonium would accumulated to concentrations high enough to inhibited cell growth. High concentration ammonium will result in high level expression of several genes concerned the central carbon and nitrogen metabolism, and thus change the metabolic mode of cells.

First, effect of nitrogen substrate to the growth of ZR1 from methane and methanol were analyzed. Nitrate salts were proved to be the best nitrogen substrate to support the growth of ZR1 from methanol and methane as carbon source. The nitrate intermediate metabolite ammonium was found to inhibit the growth of ZR1 from methanol and methane. High nitrate concentration inhibition phenomenon has long been investigated in 1991 by Park et al.(Park et al., 1991), however its inhibition mechanism in methanotrophs still lack studying. According to the substrate inhibition theory (Muchandani and Luong, 1989), the nitrate metabolic intermediate ammonium were found to accumulate, and might inhibit the growth of ZR. Kinetic study revealed that the concentration of accumulated ammonium was proportional to the original nitrate concentration in the medium. With high initiate nitrate salt, the metabolic mediates NH_4_^+^ would accumulate to more than 10 mM which was high enough to inhibit the growth of ZR1. These results indicated that worse growth of ZR1 might be owing to the disequilibrium of carbon and nitrogen metabolism. Supplying carbon skeleton to assimilate extra ammonium was proven to be a suitable strategy to relieve the inhibition effect of NH_4_^+^. The carbon skeleton replenish effect of several carbon substrate to ammonium inhibition was found to be different with methanol and methane as carbon source. Howeverα-KG was found to be the best carbon skeleton to relieve the ammonium inhibition effect. qPCR analysis indicated that, *gcvT*, *gln*A, and *ala*D genes were expressed relatively higher when ZR1 confronted with the high ammonium concentration with methane as carbon source. With addition of the C5 carbon skeletonα-KG, *gln*A gene expressed higher, and the *gcv*T gene decreased 8 times lower to ammonium accumulation condition, which indicated that, ZR1 may rely more on the third direction to assimilate extra ammonium. With methanol as carbon source, the *glt*B and *ala*D gene expressed at a relative high level at normal condition in comparison with methane as carbon source. When confronted the ammonium inhibition condition, ZR1 may decrease its *nir* and *nif* gene expression level and up-regulate the *hps* gene to further prevent the accumulation of ammonium.

In this study it was found that the nitrogen metabolic intermediate ammonium might be a signature of the nitrogen metabolism in *Methylomonas*, and induce related genes to balance the carbon and nitrogen metabolism. Besides the normal ALAD and GS/GAGOT ammonium assimilation pathway, the third Glycine synthase using the C1 carbon skeleton may also actively expressed in ZR1. Methanotrophs may utilize a complex strategy to balance the carbon and nitrogen metabolism according to the available carbon source. These findings are meaningful to reveal the complex coordination metabolic mechanism of nitrogen and carbon in methanotrophs. However, the ammonium signal transduction pathway of methanotrophs needs further study to reveal its induction mechanism.

## Acknowledgement

This work was supported by STS Project of the Chinese Academy of Sciences (No. KFJ-STS-ZDTP-065), Key project of Tianjin (No. 14ZCZDSY00157). Kind help extended by the graduate students Yanyun Guo, Mengdi Zhang, Lili Liu during the study are greatly appreciated.

